# Tardigrades exhibit robust inter-limb coordination across walking speeds

**DOI:** 10.1101/2021.03.19.436228

**Authors:** Jasmine A Nirody, Lisset A. Duran, Deborah Johnston, Daniel J. Cohen

## Abstract

Tardigrades must negotiate heterogeneous, fluctuating environments, and accordingly utilize locomotive strategies capable of dealing with variable terrain. We analyze the kinematics and inter-leg coordination of freely walking tardigrades (species: *Hypsibius dujardini*). We find that tardigrade walking replicates several key features of walking in insects despite disparities in size, skeleton, and habitat. To test the effect of environmental changes on tardigrade locomotor control circuits, we measure kinematics and inter-leg coordination during walking on two substrates of different stiffnesses. We find that the phase offset between contralateral leg pairs is flexible, while ipsilateral coordination is preserved across environmental conditions. This mirrors similar results in insects and crustaceans. We propose that these functional similarities in walking co-ordination between tardigrades and arthropods is either due to a generalized locomotor control circuit common to panarthropods, or to independent convergence onto an optimal strategy for robust multi-legged control in small animals with simple circuitry. Our results highlight the value of tardigrades as a comparative system towards understanding the mechanisms – neural and/or mechanical – underlying coordination in panarthropod locomotion.

---

The vast majority of animals need to move to survive. Tardigrades, though famed for their slow and unwieldy gait, are no exception. One of the smallest legged animals, tardigrades rely on their locomotive abilities to escape from predators, and to find food and mates [22, 26]. The first observations of tardigrades in the 18th century centered around their distinctive gait: they were described, in quick succession, as ‘water bears’ [11] and ‘Il tardigrado’ [29] due to their slow, lumbering style of walking. Beyond these initial characterizations, however, not much is known about how tardigrades move about in their environment. In more recent years, their ability to withstand environmental extremes by entering a dormant state called a ‘tun’ has garnered significant attention [3, 16, 14]. But even this famed resilience relies on locomotion: tun formation requires slow and controlled dehydration, making moving in dramatically fluctuating micro-environments an important behavioral factor in successful dehydration and rehydration [4].

Unlike other fundamental rhythmic motor programs (e.g., heartbeat or respiration), locomotion needs to be flexible and responsive to environmental stimuli. Terrestrial tardigrades in particular must traverse a complicated three-dimensional environment comprising a wide range of terrain types. Furthermore, different behavioral goals call for different walking speeds, ranging from slow exploratory walking to swift running for escape maneuvers [9]. As such, locomotor output must be tuned to both speed and substrate. Such tuning can be achieved via adjusting the kinematics of single legs (e.g., stepping frequency or step length), but often also results in changes in temporal coordination between legs.

These changes can occur in the form of transitions between discrete gaits: for instance, a horse switches from walking to trotting to galloping as it goes from slow to intermediate to high speeds [1, 2]. Alternatively, stepping patterns can lie along a continuum of inter-leg coordination patterns (ICPs). This may indicate that a single control circuit may suffice to generate all observed ICPs – that is, there need not be separate dedicated controllers for each ‘gait’. Excitingly, recent analyses have suggested the existence of such a continuum in walking insects [20, 35, 30, 8].

Are there fundamental principles behind the generation of such a continuum that can be generalized beyond insects to describe walking in other legged panarthropods? Morphological similarities in the underlying neural structure have been noted across Panarthropoda (Arthropoda + Tardigrada), supporting a sister-group relationship between these taxa [36]. However, given the large disparities in size, skeletal morphology, and environment between arthropods and tardigrades, it is unclear if these morphological parallels translate to similarities in whole-organism performance between tardigrades and arthropods.

For instance, tardigrades, with body lengths down to just several hundred microns, exist at a scale where most other organisms have opted for locomotive modes other than walking. To what extent does their scale and aquatic environment affect their inter-leg coordination and biomechanical strategy in comparison to larger legged organisms? Furthermore, little is known about how the control of locomotion in soft-bodied animals differs from those with rigid skeletons. Most soft animals are legless (e.g., nematodes or the larval stages of insects like *Drosophila*), and the lack of discrete contact points with the substrate makes it difficult to identify changes in the timing of ground interactions [32]. Deeper evaluation in Onychophorans (velvet worms) [23], and in the larval stages of holometabolous insect orders which have developed leg-like appendages called ‘prolegs’ on their abdomens (e.g., beetles [38]; sawflies, moths and butterflies [15, 31, 21]), has indicated that soft-bodied locomotion is less regular than that observed in jointed arthropods. It is unclear if this increased variability is due to differences in underlying neural connectivity, or is simply a consequence of increased degrees of freedom associated with controlling lobopodal locomotion.

Here, we utilize the framework developed from studies of insect walking to compare the biomechanical strategies used by the eutardigrade *Hypsibius dujardini* with those described in arthropods. Is walking in tardigrades described by a single continuum of ICPs? What is the relationship between the structure of the tardigrade neural system and organismal function during locomotion? We characterize the kinematics of how tardigrades generate a range of forward walking speeds, as well as how their strategies adjust given variation in environmental properties like substrate stiffness. Surprisingly, we find marked similarities in coordination between tardigrades and arthropods during spontaneous planar walking [6, 10, 20, 35, 30, 8], suggesting that either (1) there exists a common circuit for the control of forward walking in panarthropods; or (2) that tardigrades and arthropods independently converged upon a similar set of coordination strategies for locomotion despite striking differences in size, skeletal morphology, and ecology. Our findings emphasize the value of the tardigrade as a compelling comparative system to study the evolution and mechanisms of legged locomotion. Additionally, characterizing the similarities and differences between tardigrade locomotion and that of other legged organisms may also inform the development of both soft robotics and micro-scale technologies.

## Results

### Overview of tardigrade leg and claw morphology

We quantify spontaneous planar walking in the eutardigrade *Hypsibius dujardini* (see Materials and Methods). All known tardigrade species have four pairs of legs, with the fourth leg pair oriented posteriorly (Fig. 1a). Structural studies have demonstrated that while all four legs are similar in structure and show serial homologies, the number of leg muscles decreases moving anterior to posterior, with the fourth leg pair having the fewest and most divergent musculature [13]. Several kinematic parameters of the fourth leg pair show larger variation and lower dependence on walking speed relative to that in the first three pairs (Fig. S1-S3). This is in accordance with the hypothesis that the posterior legs of tardigrades are used primarily for grasping rather than propulsion in forward locomotion [25]. Despite observations of a monotonic decrease in number and monotonic increase in branching of musculature from the anterior to posterior segments in tardigrades [13], we find no significant differences in kinematic parameters or stepping precision among the first three leg pairs, suggesting that the first three leg pairs function equivalently for forward planar walking.

**Figure 1.**
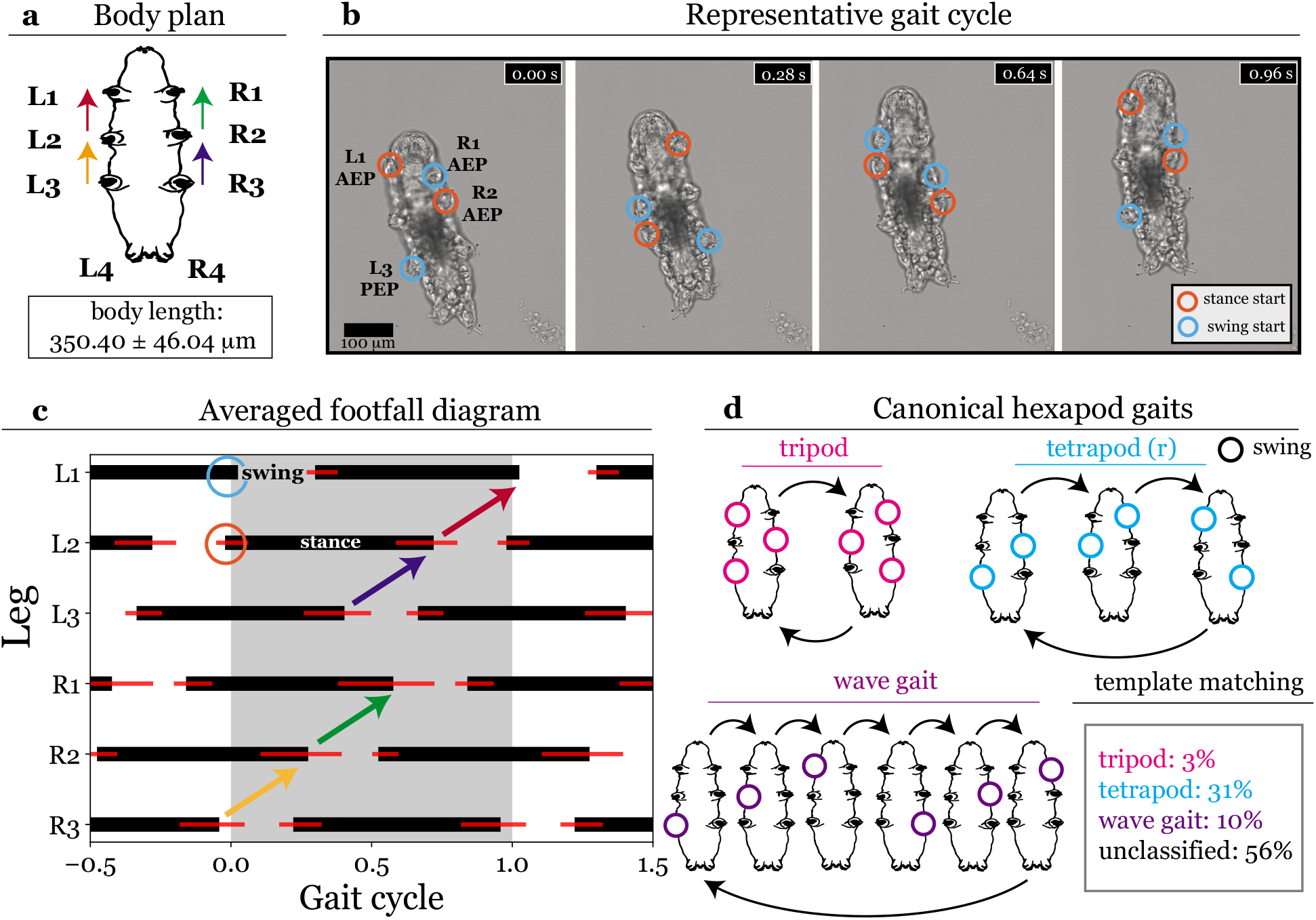
Overview of tardigrade kinematics. **(a)** Tardigrade body plan. Colored arrows denote inter-leg relationships as in the podogram shown in **(c)**. Note that kinematic data for the reduced back legs (L4, R4) are not shown here (see Supplementary Information). Mean *±* s.d. body length (measured from the tip of the nose to the attachment of the back leg pair) is provided for animals pooled between 50 kPa (*N* = 23) and 10 kPa (*N* = 20) conditions. **(b)** An exemplary stepping cycle comprising one swing and stance phase for all legs. **(c)** Podogram shows the average temporal sequence of ground contacts for legs L1-L3 (left, anterior to posterior) and R1-R3 (right, anterior to posterior). Values are normalized to cycle period of the left front leg L1, shown within the gray shaded region, [0.0 - 1.0]. Extrapolated sequences for previous [-0.5, 0) and subsequent (1, 1.5] periods are shown outside the shaded area. A total of *n* = 122 cycles (here, we define ‘cycle’ as a sequence containing one full stride from each leg) from *N* = 23 animals are shown. Mean *±* s.d. is depicted; s.d. indicated by red lines. Colored arrows highlight posterior-to-anterior propagation of ipsilateral swing events; color scheme for legs is as shown in tardigrade schematic drawing in **(a). (d)** Schematic of canonical hexapod stepping patterns: tripod, tetrapod, and wave gait. In tripod, three limbs swing simultaneously; in tetrapod, two limbs swing simultaneously; and in (pentapodal) wave gait each limb swings individually. The transitions between configurations are shown to reflect the posterior-to-anterior propagation of ipsilateral swing events (as in **(c)**). Gait template matching, a technique which classifies each stride to a ‘canonical’ pattern of phase offsets for swing initiations (see Table S1), fails to classify 56% of strides. This is largely because while the majority of strides are ‘tetrapodal’, limbs that are expected to swing simultaneously instead swing with a slight temporal offset so that anti-phase contralateral coordination is maintained.

We observe that tardigrades had considerable difficulty achieving persistent locomotion on standard, polished glass slides (Video S1). Without claw engagement, animals appear unable to successfully move forward; we find a higher probability of productive directed movement on roughened glass substrates. Hence, to more effectively capture their native environment and support claw-ground engagement, we perform all core characterizations on soft, polyacrylimide gels engineered to have a stiffness of 50 kPa where we could clearly observe claw engagement causing gel deformation (Video S2-S3; Fig. S4).

### Tardigrade kinematics across walking speeds

In our experiments, tardigrades walk with an average speed of 163.0 *±* 49.9 *µ*m/s (0.48 *±* 0.11 body lengths/s; range: 79.1 - 263.5 *µ*m/s). To initially describe the walking behavior, we examine the relationship between several kinematic parameters and speed (Fig. 2 and Fig. S1). A stride for each leg comprises a ‘swing’ phase, in which the leg is lifted and takes a step, and a ‘stance’ phase, in which the leg is in contact with the ground. Step amplitude, defined as the distance between the posterior extreme position (PEP) measured at lift-off of a leg at the start of a swing and the anterior extreme position (AEP) measured at touch-down of the same leg at the end of swing, increases with forward walking speed (Fig. 2a). Stride period decreases with walking speed, plateauing at walking speeds of approximately *v >* 100*µ*m/s (Fig. 2b). As speed increases, stance duration is modulated strongly while swing duration remains relatively constant (Fig. 2c). Both stance duration and stride period show a hyperbolic relationship with speed, as observed in insects (*Drosophila* [35]; stick insect *Carausius morosus* [7]; desert ant *Cataglyphyis fortis* [33]). In line with studies in arthropods, we find that swing duration is coordinated with stride period (*ρ* = 0.54, *p <* 0.001).

**Figure 2.**
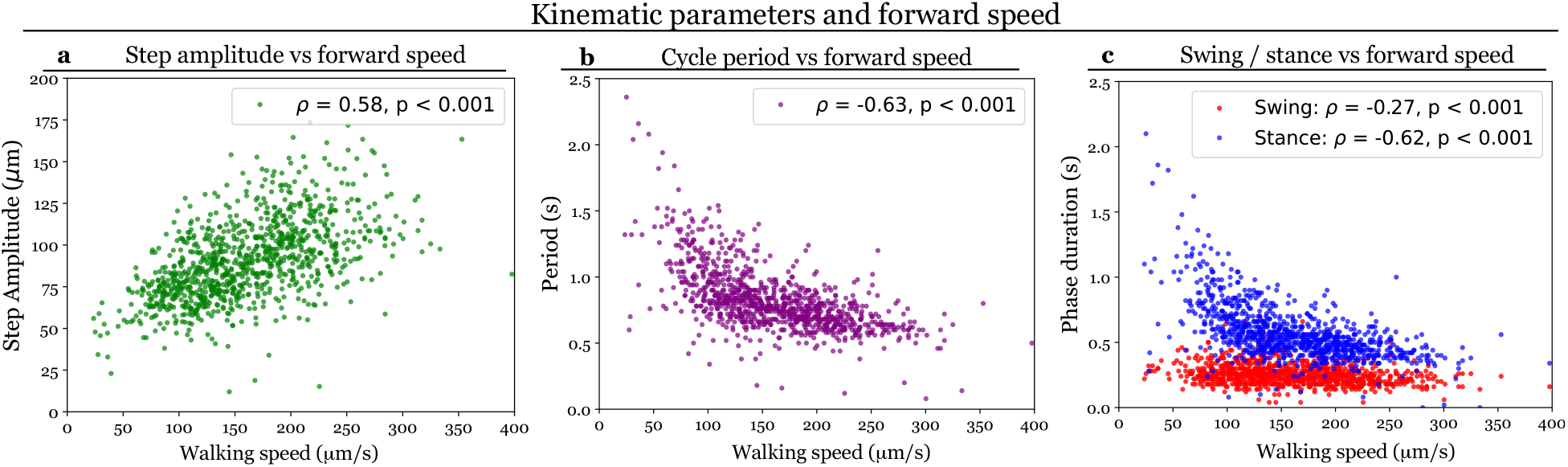
Leg stepping parameters relative to forward speed. Each point represents a stride (stance + swing) for an individual leg, *n* = 989 strides are shown. Data for the first three leg pairs are pooled, individual fits for each leg pair are provided in Figure S1. Stride length **(a)** smoothly increases and period **(b)** smoothly decreases with walking speed, suggesting that tardigrades modulate both stride length and stepping frequency to increase forward speed. **(c)** Each step is composed of a swing (leg lifted) and stance (leg on the ground) period; swing duration stays roughly constant with speed, while stance duration is modulated, decreasing with increasing speed.

This relative modulation is cleanly characterized by changes in the duty factor, the proportion of a gait cycle spent in stance phase. Duty factor changes smoothly with walking speed across the majority of arthropods, which is consistent with observations that arthropod stepping patterns lie along a continuum. Tardigrades, like all arthropod species surveyed, show a smooth relationship between duty factor and forward walking speed (Fig. 3), suggesting that they similarly do not display discrete gaits but continuously transition between ICPs. Faster walking speeds generally result in stepping patterns with lower duty factors – i.e., legs spend proportionally less time on the ground the faster the animal is moving. Species that do not utilize a wide range of stepping patterns therefore do not show significant changes in duty factor over observed speeds. For instance, the adult stick insect *C. morosus* is a slow, careful walker and overwhelmingly favors the stable tetrapodal coordination (in which four legs are kept on the ground at any given time), maintaining a near-constant duty factor across its small range of natural walking speeds (Fig. 3 inset, purple). The tardigrade *H. dujardini*, which has to navigate similarly complex terrain (albeit at a very different length scale), displays a relatively weak correlation between duty factor and speed in (*ρ* = *−*0.33, *p <* 0.001). This may suggest that living in and moving through variable environments results in a preference for a consistent, stable stepping pattern over the ability to walk at a wide range of speeds.

**Figure 3.**
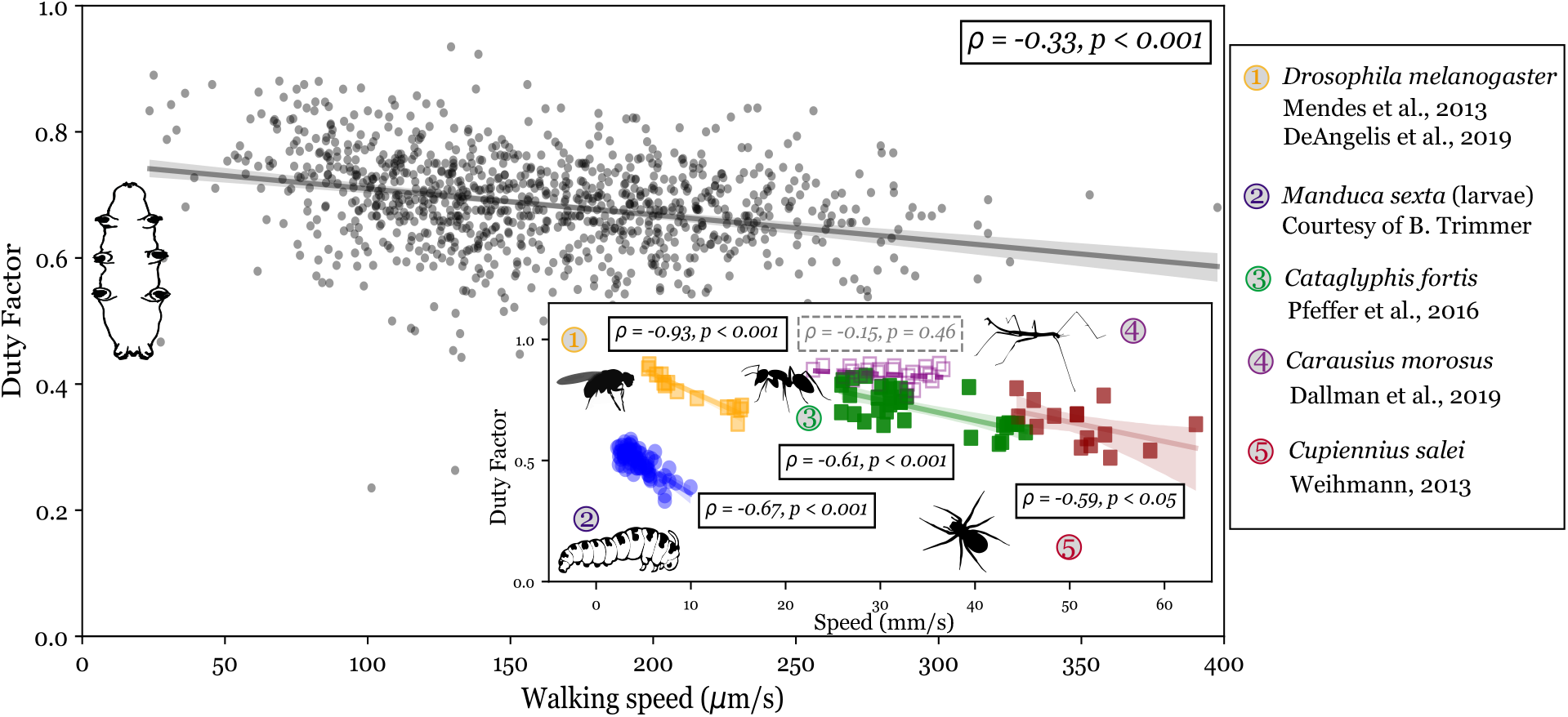
Comparison of duty factor vs forward speed across Panarthropoda. Tardigrades maintain a relatively constant duty factor across all observed forward walking speeds; we find a weak negative correlation between duty factor and walking speed (*ρ* = −0.33, *p <* 0.001). As in Fig. 2, each point represents a stride for a single leg; a total of *n* = 989 strides are shown. Inset figure shows the duty factor vs walking speed relationship for several arthropod species during slow walking. All species except the stick insect (inset gray points, data from [7]) show significant inverse relationships between forward speed and duty factor. Linear regression fits are shown as solid lines alongside 95% confidence intervals; fits for which *p >* 0.05 are shown as dotted lines.

### Smooth transitions between stepping patterns

As previously noted, stepping patterns in hexapods are often grouped into three canonical ‘gaits’ (Fig. 1d). In tripod gait, two sets of three limbs swing together; in tetrapod gait, three groups of two limbs swing together; in wave gait, each limb swings alone [35]. Our experiments show that *H. dujardini* primarily prefer a tetrapodal stepping pattern across speeds; an exemplary gait cycle is shown in (Fig. 1b). This is consistent with our finding that tardigrade duty factor maintains the expected value for tetrapodal coordination (*df* = 0.67) at all observed speeds (Fig. 3).

As previously mentioned, there is mounting evidence that stepping patterns in insects (and perhaps, more generally) do not correspond to distinct gaits, but instead form a speed-dependent continuum of inter-leg coordination patterns (ICPs) [8, 30, 35]. Tardigrades, similarly, do not adhere cleanly prescribed canonical gaits. Classification using gait template matching results in 56% of tardigrade strides remaining ‘unclassified’ and non-canonical, even after allowing for behavioral variance and tracking discrepancies (Fig. 1d). In accordance with our initial observations, approximately 70% of classified strides are sufficiently aligned with ‘idealized’ tetrapod coordination (Fig. 1d; see Materials and Methods); this alignment increases with walking speed (Fig. S5).

To determine how this spectrum of gaits arises, we considered that studies in *Drosophila* have indicated that tuning a single parameter, stance duration, can generate the spectrum of observed walking patterns [8]. Here, we find that tardigrades show a smooth, continuous relationship in both stance duration (Fig. 2c) and duty factor (Fig. 3) with speed, supporting our hypothesized lack of distinct gaits, and lack of discrete transitions between them.

### Simple, local coordination rules explain tardigrade ICPs

To better understand what kind of controller is responsible for tardigrade locomotion patterns, we turn to studies demonstrating that a small set of simple ‘coordination rules’ is sufficient to generate the continuum of observed insect ICPs during planar walking [24]. These locally-distributed rules describe how a leg affects the likelihood of the initiation of a swing event in an anterior or contralateral neighboring leg [6, 10]. *Rule 1* states that a leg’s stance-to-swing transition is suppressed while its neighbor is in swing, while *Rule 2* states that the likelihood of lift-off increases once the neighboring leg touches down. Both rules have been shown to be stronger between ipsilateral rather than between contralateral leg pairs [10, 24].

A unifying hypothesis for the observed continuum of stepping patterns [8] and decentralized control of insect walking [10, 24] is rooted in the structure of the arthropod nervous system. As anatomical studies have highlighted the similarities between the tardigrade and arthropod nervous systems [19, 36], we hypothesize that stepping patterns in tardigrades should exemplify several key attributes of arthropod ICPs. Each pair of legs – front, middle, and hind leg pairs in arthropods – is controlled by its own ganglion of the ventral nerve cord (VNC), each of which consists of two right and left hemi-ganglia, which control the corresponding legs. The three segmental ganglia are linked by longitudinal connectives [19]. A simple hypothesis consists of mutual inhibitory coupling between each right and left hemi-ganglion and a posterior-to-anterior inhibitory coupling between the longitudinal commissures connecting ipsilateral ganglia [8]. This is analogous to Rule 1 (Fig. 4a and Fig. S6c). Indeed, our analysis of inter-leg coordination compellingly suggests that both Rule 1 and Rule 2 are active between ipsilateral leg pairs in tardigrades. In accordance with Rule 1, our data show that the likelihood of a swing initiation is nearly zero after its posterior ipsilateral neighbor lifts off (Fig. 4a and Fig. S6c). This likelihood sharply rises after the time since its posterior ipsilateral neighbor’s swing initiation surpasses the sample average swing duration *(t*_swing_) = 0.18 (normalized to one cycle length). To check compliance with Rule 2, we examine the likelihood of a leg lifting off into swing phase after its posterior ipsilateral neighbor completes its swing and touches down (Fig. 4b). As predicted, we find that this probability rises sharply immediately after touch down of the ipsilaterally posterior leg.

**Figure 4.**
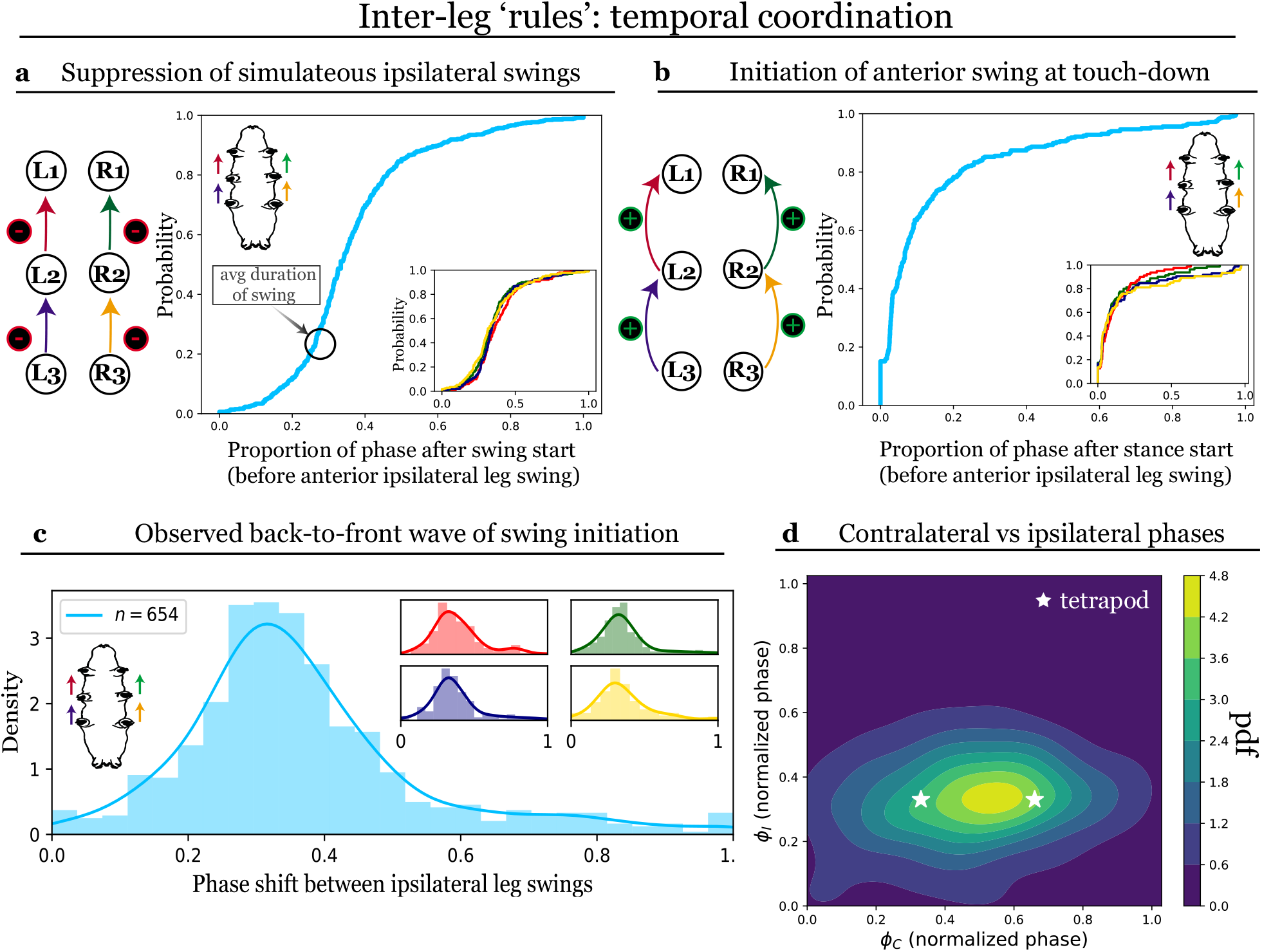
Temporal coordination between ipsilateral leg pairs. Plots represent data pooled from all leg pairs; a total of *n* = 654 strides are shown. Insets show pairwise inter-leg relationships, with color scheme as demarcated by the arrows on tardigrade schematics. Cumulative distribution functions (CDFs) show that leg swings are **(a)** suppressed immediately following the swing initiation of the posterior ipsilateral neighbor and **(b)** initiated after the posterior ipsilateral leg has touched down into stance. **(c)** Histogram and probability density of observed phase shift between ipsilateral legs show that tardigrades maintain a posterior-to-anterior wave with a phase difference 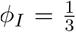 between swing onsets. Similar observations have been made in several arthropods [6, 35, 30]. Pairwise comparison using the Kolmogorov-Smirnov test found no significant differences between any leg pairs, after controlling for multiple testing. **(d)** Joint distribution of the phase difference between contralateral (*Φ*_*C*_) and ipsilateral (*Φ*_*I*_) leg pairs. Preferred coordination shows characteristic phase differences different from those observed in ‘idealized’ tetrapod. Notably, while tardigrades prefer the expected 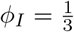 contralateral leg pairs display anti-phase coordination 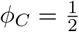

Compliance with these rules in tardigrades results in a back-to-front wave of swing initiations across walking speeds (Fig. 1c), a pattern observed in a broad range of arthropod taxa [6, 10, 20, 35, 30, 8]. The average ipsilateral offset in our data, *Φ*_*I*_ = 0.36 *±* 0.17, is approximately as predicted for tetrapod leg coordination patterns (ipsilateral phase offset for ‘ideal’ tetrapod: 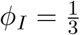 Phase offsets are equivalent between all ipsilateral leg pairs (Fig. 4c), suggesting a similar role for all three sets of legs in forward planar walking. Leg pairs maintain this offset across walking speeds; coordination is more variable at very low speeds, as was found in previous studies [35, 8].

Inter-leg coupling in the context of these rules is significantly weaker (if active at all) between contralateral leg pairs (Fig. S6c). Our data finds that all contralateral leg pairs show an average anti-phase preference. Such a value may arise from a bimodal distribution with twin peaks at 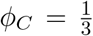 and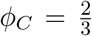, as might be expected from the two idealized tetrapod patterns (Fig. 1d). However, we find contralateral phase differences to be normally distributed about 0.5 (Fig. 4d and Fig. S6b). This relationship arises from a slight cross-body offset in swing initiations, such that limbs that would be expected to swing simultaneously are slightly offset in their liftoff time. Interestingly, previous work in *Drosophila* has also reported anti-phase contralateral coordination across forward walking speeds [35, 30, 8]; though it has not yet been explicitly tested, this pattern may be common to tetrapod-like stepping patterns across arthropods.

### Changes in limb kinematics and ICPs with substrate stiffness

Natural terrain is rarely homogeneous, and legged animals often need to cope with environmental inconsistencies such as changes in substrate roughness [5, 34] or stiffness [21]. These irregularities often constrain organismal performance, and can result in constraints on walking speed as well as changes in limb kinematics. Limnoterrestrial tardigrades, like *H. dujardini*, walk along soft, uneven plant matter. Having established earlier that claw engagement is essential for *H. dujardini* and that hard, flat surfaces posed a particular challenge, we next explore the importance of substrate stiffness, hypothesizing that particularly soft substrates may pose problems to proper claw engagement[18].

To assess the role of stiffness, we compare tardigrade locomotion on our standard, 50 kPa gels to that on 10 kPa gels (see Materials and Methods). Tardigrade walking speed on the soft gels decreases nearly two fold relative to performance on the stiff gels (160.5 *±* 57.8*µ*m/s on the 50 kPa substrate vs 91.0 *±* 32.0*µ*m/s on the 10 kPa substrate; see Fig. S4). Because tardigrades do not always walk at steady speeds, we compare walking speed distributions on 50 kPa and 10 kPa substrates to rule out the apparent slower speed coming from an increase in stop-start motion. However, speeds on both substrates are distributed unimodally, suggesting that the change in walking speed with substrate stiffness is due to a shift towards lower preferred walking speeds rather than more frequent acceleration and deceleration. More specifically, the observed reduction in speed is achieved largely through changes in stride period rather than step length; within each stride, the stance duration varies significantly between conditions, while swing duration does not

(Fig. 5a). As such, higher duty factors are associated with stepping patterns on the 10 kPa substrate (Table S2), perhaps indicating greater locomotive effort necessitating longer ground contact times.

**Figure 5.**
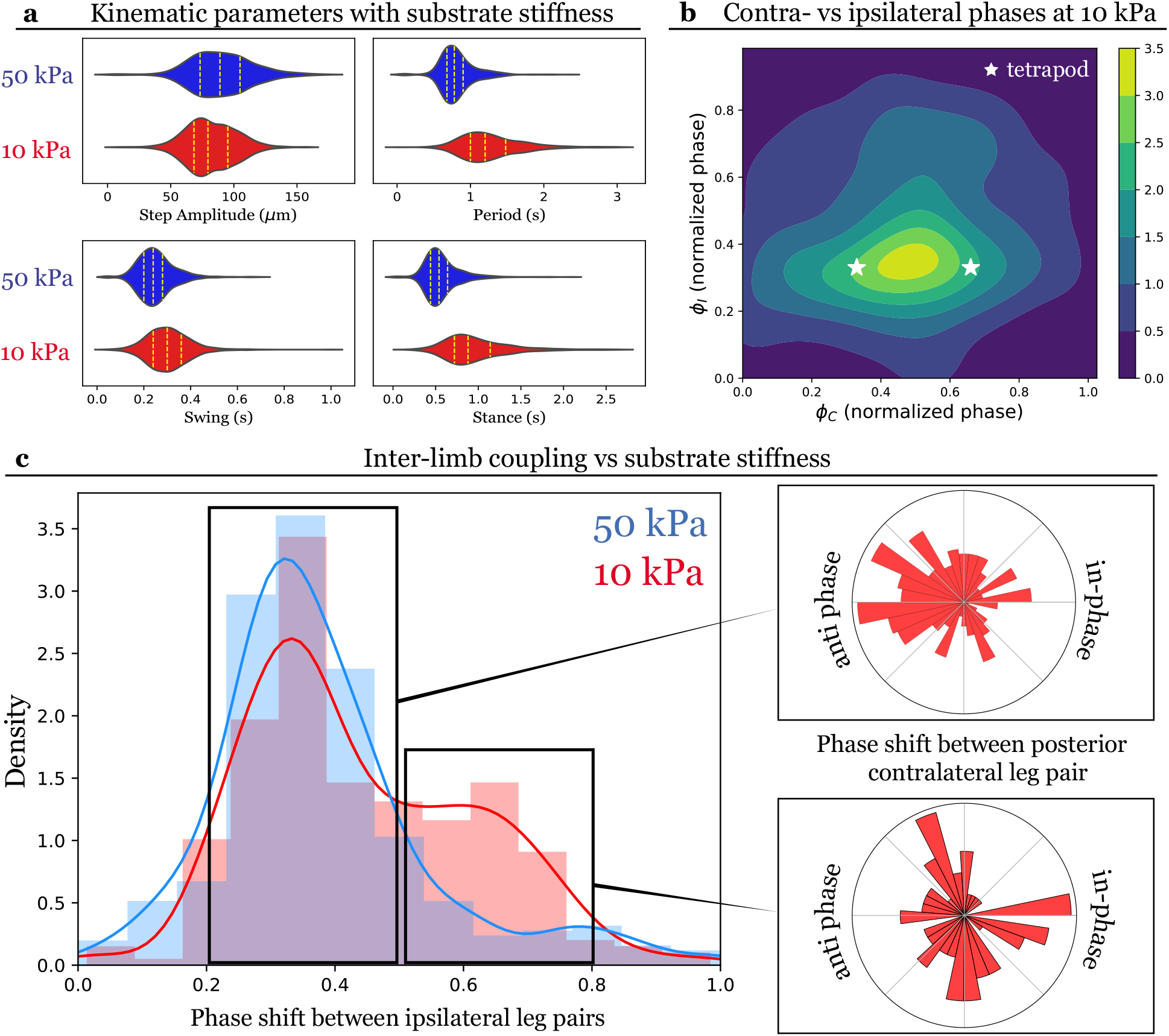
Effects of substrate stiffness on walking kinematics and inter-leg coordination. **(a)** Tardigrades adjust period (top right panel) and stance duration (bottom right panel) with changes in substrate stiffness. Distributions for step amplitude (top left panel) and swing duration (bottom left panel) are largely unchanged between stiffness conditions. **(b)** Joint distribution of the phase difference between contralateral (*Φ*_*C*_) and ipsilateral (*Φ*_*I*_) leg pairs on the 10 kPa substrate. As on the 50 kPa substrate, tardigrades prefer the expected tetrapodal ipsilateral phase difference 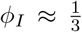 but display anti-phase contralateral coordination 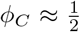. Larger variance in ipsilateral phase is apparent than on the 50 kPa substrate. **(c)** Comparison of the marginal ipsilateral phase difference distributions on 50 kPa and 10 kPa substrate shows an additional peak at 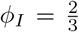 on the 10 kPa substrate (red). This arises because of transient in-phase stepping by the neighboring posterior leg pair (bottom inset), deviating from the preferred anti-phase (top inset).

We also explore how substrate stiffness modulated specific inter-leg control rules, and find that inter-leg phase relationships are mostly robust to changes in substrate stiffness (Fig. 5b). The preferred stepping patterns on the 10 kPa substrate maintain several key features to that observed on the 50 kPa substrate: (1) ipsilateral swing events proceed in a posterior-to-anterior fashion; (2) adjacent ipsilateral legs show a preferred phase difference 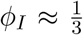 and (3) contralateral leg pairs show a preference for anti-phase coordination 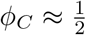 However, we find a smaller second peak in the distribution of ipsilateral phase differences at 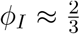 of strides taken on the 10 kPa substrate (Fig. 5b,c). Closer analysis reveals that this second peak arises due to the posterior-most leg pair showing in-phase rather than anti-phase coordination (Fig. 5c). These alternate stepping patterns arise when a leg pair steps in phase (*Φ*_*C*_ = 0), and consequently, the leg pair in front of it may either adopt an (1) in-phase contralateral coordination to maintain a constant ipsilateral phase difference 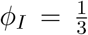, or an (2) anti-phase contralateral coordination to ‘reset’ to the preferred tetrapod-like pattern, which leads to one side showing an ipsilateral difference of 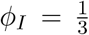 and the other side showing a phase difference 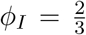. The latter strategy is preferred, with in-phase contralateral stepping patterns being largely transient; however, we did observe rare instances of a sustained ‘galloping’ gait as would result from the former strategy (Video S4).

## Discussion

In addition to their famed resilience under conditions of extreme stress, tardigrades also display a remarkable robustness in their day-to-day activities. Tardigrade morphology is strongly conserved across species that live in and move through a large range of habitats, including marine, freshwater, and limnoterrestrial environments. Here, we characterize how kinematics and inter-leg coordination in the limnoterrestrial tardigrade *H. dujardini* adapt to changes in walking speed and substrate properties. We find that tardigrade stepping patterns change smoothly with walking speed, rather than displaying sharp transitions between discrete ‘gaits’. Further, the observed patterns reproduce key features in the spectrum of ICPs characterized in various insect species, namely: (1) a posterior-to-anterior wave of ipsilateral swing initiations across all walking speeds; and (2) a general preference for anti-phase coordination between contralateral legs. More generally, we find that the observed spectrum of tardigrade inter-leg coordination patterns emerges naturally from surprisingly small set of local coordination rules derived from behavioral observations in insect species [10].

These functional parallels are particularly striking given the large disparities in size, skeletal morphology, and environmental between tardigrades and insects. Tardigrades are approximately 500,000 times smaller than stick insects, and yet ‘rules’ derived from behavioral studies in *C. morusus* sufficiently and accurately describe walking patterns in *H. dujardini* [6, 10]. Whether these commonalities arise from shared ancestral structures remains an exciting open question [36, 19, 27].

One hypothesis points to anatomic similarities in underlying neural circuitry. A simple model for locomotor control in insects was proposed based on studies in walking *Drosophila*, and is built upon the anatomy of the ventral nerve cord (VNC). This model supposes a posterior-to-anterior inhibitory coupling between ipsilateral neuropil, as well as mutual inhibitory coupling between contralateral neuropil of the VNC [8]. There are various striking similarities in tardigrade and arthropod VNC structure: (1) the VNC comprises three segmented ganglia, each associated with a walking leg pair; and (2) each ganglion is divided into left and right hemiganglia linked by contralateral projections. In contrast, the closely-related velvet worms – which, like tardigrades, are among the only extant soft-bodied walkers – have two laterally-located non-segmented ganglia [36, 19]. This disparity in underlying neuroanatomy may account for the differences between onychophoran gaits and those observed in tardigrades and arthropods [23]. As such, comparative analyses making use of VNC structure indicate a sister-grouping between tardigrades and arthropods [19, 36].

It is an intriguing hypothesis that there may exist a common simple locomotor circuit underlying walking in panarthropod species, which has been modified along certain clades due to specific pressures on organismal performance (e.g., in-phase contralateral coordination in crayfish swimmerets [37]). An example of one such modification is the modulation of inter- or intra-segmental connections in response to changes in the environment, e.g., varying surface stiffness. The behavioral flexibility observed in arthropod walking is partially ascribed to the modular nature of segmented body plans. Varying of substrate stiffness is likely salient for tardigrades because they lack a rigid skeleton, and accordingly may utilize a biomechanical strategy distinct from most adult insects that relies crucially on environmental stiffness [18]. Analyses in the soft-bodied hawk moth larvae found that *Manduca sexta* caterpillars did indeed sense changes in substrate stiffness and adjusted their stepping patterns to accommodate these changes [21]. Studies in several insect species suggest that ipsilateral (inter-segmental) synchronization dominates inter-leg coupling patterns, while contralateral (intra-segmental) coupling is adjustable [34, 28, 24]. Consistent with these observations, tardigrades show changes in their contralateral phase offset but largely preserve ipsilateral coordination on soft substrates. We observe both transient strides in which contralateral limbs step in phase, as well as occasional sustained in-phase contralateral coordination resulting in a ‘bounding’ or ‘galloping’ gait. Interestingly, several species of dung beetle in the genus *Pachysoma* have also been observed to maintain a galloping gait across shifting sands [28].

However, alternative analyses – including molecular analyses [17] and comparative studies of brain structure [27] – group onychophorans together with arthropods, an interpretation which suggests that functional analogues between tardigrade and arthropod walking have independently evolved. In this case, the similarities in underlying circuitry controlling tardigrade and insect walking may not be ancestral. It may be that the observed set of coordination patterns, which only require a single simple controller, might be preferable in small animals with small circuits for limb control [8]. Previous studies have also proposed that the magnitude and robustness of static stability during walking may affect preference for certain ICPs [30]. It is unclear, however, whether stability or the need to stay upright would be a concern for tardigrades, which are far smaller than stick insects or even fruit flies, and walk underwater with the assistance of buoyancy. Parallel convergence onto similar inter-leg coordination strategies by tardigrades and arthropods is intriguing given their varied ecology, disparities in size, and difference in skeletal structure between the two groups, and can provide significant insight into general design principles for efficient and robust control of multi-legged locomotion.

A more definitive distinction between these scenarios will require deeper functional studies combined with molecular, phylogenetic, and anatomical analyses. We note that this work focuses on analyzing spontaneous walking in *H. dujardini*, and that our data is accordingly constrained both in number of species considered and in the range of walking speeds observed. Future work should expand upon both of these parameters, both by surveying walking dynamics across Tardigrada (keeping in mind the wide ecological range of the phylum) and by perturbation or manipulation experiments that may expand the observed range of walking speeds.

Tardigrade walking also poses several fundamental mechanistic questions. Tardigrades are among the smallest legged animals, and, given their ecological success, investigations into their biomechanical strategy provide valuable insight into the scaling of efficient polypedal walking in various ecological conditions. For instance, the common preference for tetrapod-like coordination in both tardigrades and far larger species like the stick insect points to the selective importance of static stability in species that regularly navigate variable, three-dimensional terrain [12, 10]. Similarly, the shift towards in-phase contralateral stepping on soft substrates in tardigrades is mirrored in the evolved ‘galloping’ gait of desert-dwelling beetles orders of magnitude larger in size [28]. This common strategy may reflect an energetic benefit to in-phase contralateral coordination on unstable or shifting terrain that holds across a remarkably large range of length scales. Furthermore, tardigrades are one of the only soft-bodied animals that walk using ‘true’ legs, which allows for cleaner characterization of coordination and kinematic strategy than is possible in soft animals that crawl without discrete ground contacts. To this end, our findings here highlight the value of tardigrades both as a comparative system towards understanding the mechanisms underlying coordination in panarthropod locomotion and as an organismal system uniquely positioned to inform our understanding of the design and control of small, soft-bodied locomotive systems, from organisms to robots.

## Supporting information

Supplemental Video 1. Tardigrade walking on smooth glass

Supplemental Video 2. Tardigrade walking on gel of 50 kPa stiffness.

Supplemental Video 3. Tardigrade walking on 50 kPa gel, sagittal view.

Supplemental Video 4. Tardigrade walking on gel of 10 kPa stiffness.

## Acknowledgments

We thank Barry Trimmer for data on *Manduca sexta* larva kinematics and many helpful discussions.

J.A.N. was supported by a James S. McDonnell Foundation Fellowship for Studying Complex Systems, and Fellowships from All Souls College at the University of Oxford and the Center for Studies in Physics and Biology at the Rockefeller University. D.J.T. thanks Princeton for a SEAS Innovation Grant from the Helen Shipley Hunt Fund. L.A.D. was supported by the Jonas Salk Award.

