## Supplementary Text, Figures, Tables. for "Tardigrades exhibit robust inter-limb coordination across walking speeds"

**2 Supplementary Tables** **5**

**3 Supplementary Figures** **7**

### 1 Materials and Methods

#### 1.1 Tardigrade Husbandry

Specimens of the eutardigrade *Hypsibius dujardini* were purchased from Carolina Biological Supply. Tardigrades were maintained in Chalkey's medium [3]. Animals were fed *Chlorococcum* sp. algae; algae was purchased from Caroline Biological Supply and maintained in Bold's Basal Medium (BBM). Cultures were maintained at 20-25°C under a 16 hr light/8 hr dark cycle.

#### 1.2 Experimental procedure and imaging

We studied tardigrades walking on polyacrylamide gels of stiffnesses approximately 50 kPa and 10 kPa. An initial gel precursor solution (40% acrylamide, 2% bis-acrylamide) were diluted in 1mL of MiliQ purified water according to measurements made in [2]: 8%/0.4% acrylamide/bis-acrylamide for 50 kPa gel substrate and 8%/0.1% acrylamide/bis-acrylamide for 10 kPa gel substrate. Solutions were degassed before gelation. To initiate gelation, 5$\mu$L of 100mg/mL ammonium persulfate was added to 1 mL of diluted precursor solution, followed by 0.5$\mu$L of N,N,N',N'-Tetramethyl-ethylenediamine accelerator (TEMED). After being mixed by gentle pipetting, 20 $\mu$L of the gel solution was added to a glass bottom dish and flattened with an 18 mm round coverslip. Glass bottom dishes were treated with Bind-Saline solution, washed with ethanol, and air dried before addition of solution. Gels were polymerized at room temperature (23°C) for 30 minutes, after which the coverslip was removed carefully from the gel surface using tweezers. Further details on preparation and stiffness measurements of polyacrylamide gel substrate is provided in [2].

Using a pipette, 3 mL of tardigrade-containing solution was removed from culture. This solution was filtered using 100$\mu$m mesh to remove algae, and 400 $\mu$L of filtered tardigrade culture was added onto the gel substrate for imaging. Tardigrades were allowed to settle for 10 minutes, after which an additional 3 mL of spring water was added to assure tardigrades were kept submerged throughout imaging. Animals were imaged using differential interface contrast (DIC) microscopy on an inverted microscope (Nikon Ti2, 20x/0.75 objective; or Zeiss Primovert, 10X/0.3 for locomotion on pure glass surfaces). Videos from the ventral view of the animal were recorded at 60 fps using a Basler acA2040-90um monochrome CMOS camera. Several videos were obtained for each animal, such that 5-10 complete strides for each tardigrade could be extracted.

#### 1.3 Kinematic and statistical analysis

Tardigrades were recorded walking spontaneously across gel substrates. We recorded trials of straight walking containing at least 2 complete cycles per leg. For each animal, between 5-15 complete cycles were recorded ($N = 23$ animals on 50 kPa gel substrate, $N = 20$ animals on 10 kPa gel substrate, and $N = 4$ animals on glass). Recorded videos from the ventral view were then evaluated frame-by-frame semi-automatcally.

Exact time and location of leg lift-off (swing initiation) and touch-down (stance initiation) events, as well as frame-by-frame position of head, center of mass (COM), and tail were visually determined and tracked using the ManualTracking plugin in ImageJ [6]. Positions for head, COM, and tail were defined by the tip of the tardigrade's nose, the midpoint between the second leg pair, and the point of attachment of the back leg pair to the tardigrade's body, respectively. Tracking of leg kinematics was only done for videos taken on gel substrates; only COM position with time was recorded for glass trials. All measurements were made in ground-fixed coordinates. Data obtained in this manner were then processed using Python. All raw and processed data, as well as analysis code, will be available at https://github.com/jnirody/waterbears.

A stride period for an individual leg was defined as the time difference between two consecutive lift-off events. Each stride comprises a stance and a swing. The durations of swing and stance were calculated as the time difference between a lift-off event and the subsequent touch-down (swing) or the time difference between a touch-down and the subsequent lift-off (swing). Temporal coordination of leg pairs was determined using the relative timings of swing onsets.

Step amplitude for an individual leg was determined as distance between the point of liftoff (posterior extreme position, PEP) and the location of the subsequent touch-down (anterior extreme position, AEP). As in [10], we use step amplitude instead of stride length, which is calculated instead as the distance between two consecutive AEPs. This is because stride length is more affected than step amplitude by factors unrelated to active changes by the animal (e.g., slipping or the effects of a softer substrate). In contrast, step amplitude is more tied to an active change in the tardigrade's kinematics.

Walking speed was calculated as the change in the measured position of the center of mass (COM). The walking speed associated with a particular stride (e.g., as in Fig. 2 or Fig. S4b,c) was calculated as follows: we first determined the relevant time period (in the case of a stride period, this would be the time difference between two lift-off events). We then used the displacement of the COM between these two points to calculate an averaged speed over that interval. If walking speed was considered alone (e.g., as in Fig. S4a), speed was averaged in the same manner over non-overlapping 60-frame (1 second) windows.

Correlation coefficients between variables were calculated over the entire observed walking speed range; we used Spearman $\rho$ to determine correlation due to the non-linear relationships between several of our kinematic variables. Data in most plots presented were pooled across walking legs (the first three leg pairs). Leg-separated data was presented in Fig. S1. Kolmogorov-Smirov tests showed no significant differences in any kinematic parameter between legs after correction for multiple testing. Regression fit lines (e.g., in Fig. S1) and density distribution fits to histogram data (e.g., in Fig. 5c and Fig. S2) were computed using Python package seaborn [9] and are intended to guide the eye. Joint distributions of phase angles (e.g., Fig. 3d, Fig. 4b, and Fig. S6) were computed using a kernel density estimate using Gaussian kernels; we note that this may have resulted in slight over-smoothing for the $\phi_C$-$\phi_I$ diagram on the 10 kPa substrate (Fig. 5b). To this end, the marginal distribution for ipsilateral phase differences is provided in Fig. 5c. All fits were computed using Python package scipy stats [8].

#### 1.4 Comparison of duty factor vs speed relationship across arthropod species

Duty factor is defined as the proportion of time spent in stance phase; specifically, for each stride, it was calculated as the ratio of the stance duration to the stride period. All statistics were computed as described above for other kinematic parameters. Duty factors for other arthopod species were extracted from published articles as cited. For some articles, tabular data was not available; in these cases, data was extracted from paper figures using the R package digitize [5]. Kinematic data for *Manduca sexta* caterpillars was provided courtesy of Barry Trimmer. All numerical data used to make Fig. 3 is available at https://github.com/jnirody/waterbears.

#### 1.5 Inter-leg coordination and gait template matching

For gait template matching, we utilize a similar framework to that described in [4]. A gait was assigned to a block of time in which each walking leg completed one full stride; the boundaries of this period are from the earliest swing onset to the latest stance onset. To determine the assigned gait, phase relationships were calculated as the onset of swing relative to the stepping period of the right hind leg (R3). Table S1 shows the idealized step pattern for each canonical gait; note that 'tetrapod' actually comprises two mirror image coordination patterns, denoted as 'tetrapod 1' and 'tetrapod 2'. Because perfectly synchronous swing movements were exceedingly rare in our data, and to account for human error in tracking, we tolerate a deviation from ideal swing relationships by $\pm 0.12$. When assigning a gait, we allowed one erroneous step for a single leg per full cycle. This is equivalent to the allowances given to stick insects in [4].

Though we use the terminology often seen in the literature referring to the described canonical gaits for hexapodal walking, we note that this is purely for descriptive purposes. Insect (and likely panarthropod) walking is better represented by a continuum of inter-leg coordination patterns rather than by distinct, discrete gaits. This is emphasized in our results by the fact that 56% of strides were unable to be classified into one of these three categories using the generous gait template matching framework described above.

Therefore, in order to more accurately describe the walking patterns observed during our experiments, we focused on measuring pairwise inter-leg coordination, particularly between ipsilateral and contralateral neighbors. We define an ipsilateral leg pair as two neighboring legs on the same side (e.g., R1 and R2), and contralateral leg pairs as two legs directly opposite each other (e.g., L2 and R2). Phase differences between leg pairs are denoted throughout as $\phi_I$ between ipsilateral leg pairs and $\phi_C$ between contralateral leg pairs. The leg within the pair which swings first within a full cycle (comprising swing events of all six legs, as described above) is considered the reference leg; the phase offset is normalized with respect to its stride period. For example, let us consider the ipsilateral leg pairing (R1,R2); in our data the posterior leg is always the reference leg in ipsilateral leg pairings. Consecutive swing initiations of R2 demarcate the boundaries of the period $(t_0, t_1]$. Then if R1 swings at time $t_s$, the phase difference $\phi_I^{R2-R1}$ is given by:

$$\phi_I^{R2-R1} = \frac{(t_s - t_0)}{(t_1 - t_0)}. \tag{1}$$

Phase differences between contralateral leg pairs are calculated equivalently.

#### 2 Supplementary Tables

| Canonical gait | R3 | L3 | R2 | L2 | R1 | L1 |
|---|---|---|---|---|---|---|
| Tripod | 0 | 1/2 | 1/2 | 0 | 0 | 1/2 |
| Tetrapod1 | 0 | 2/3 | 1/3 | 0 | 2/3 | 1/3 |
| Tetrapod2 | 0 | 1/3 | 1/3 | 2/3 | 2/3 | 0 |
| Wave | 0 | 3/6 | 1/6 | 4/6 | 2/6 | 5/6 |

Table S1: Phase relationships between legs in canonical hexpod gaits; relative to swing start of R3.

| Parameter | 50 kPa (mean ± sd) | 10 kPa (mean ± sd) |
|---|---|---|
| Number of animals | 23 | 20 |
| Body length ($\mu$m) | $338.12 \pm 45.04$ | $364.53 \pm 44.09$ |
| Walking speed ($\mu$m/s) | $160.50 \pm 57.79$ | $91.0 \pm 32.0$ |
| Step length ($\mu$m) | $90.52 \pm 23.70$ | $82.07 \pm 19.74$ |
| Period (s) | $0.82 \pm 0.24$ | $1.27 \pm 0.41$ |
| Stance duration (s) | $0.57 \pm 0.21$ | $0.97 \pm 0.37$ |
| Swing duration (s) | $0.25 \pm 0.08$ | $0.30 \pm 0.09$ |
| Duty Factor (%) | $0.68 \pm 0.08$ | $0.75 \pm 0.07$ |

Table S2: Mean ± sd for kinematic parameters on 50kPa and 10kPa substrates.

### 3 Supplementary Figures

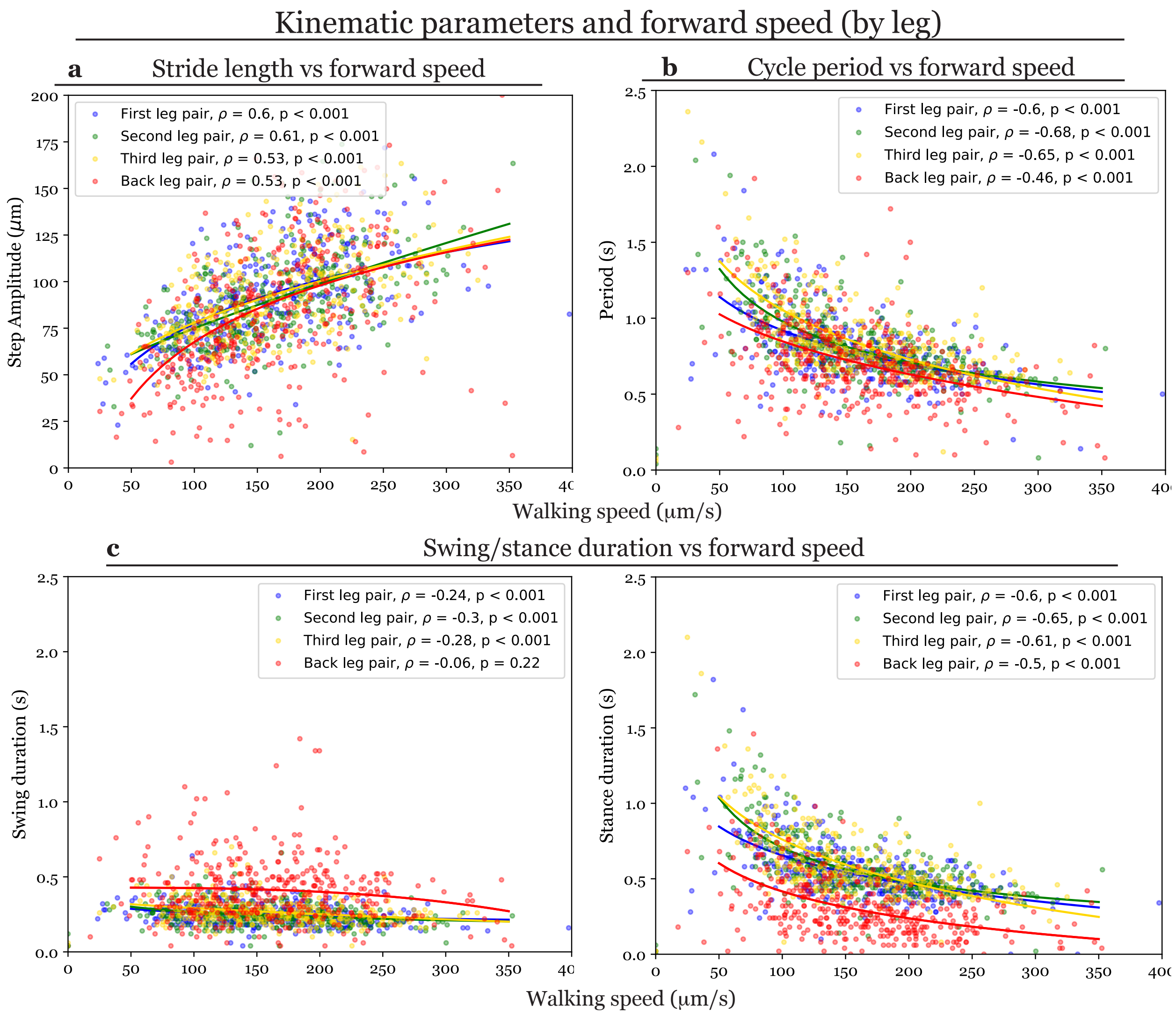

Figure S1: Relationship between forward walking speed and **(a)** step amplitude, **(b)** period, and **(c)** swing and **(d)** stance duration for each leg pair. Fit lines are provided to guide the eye.

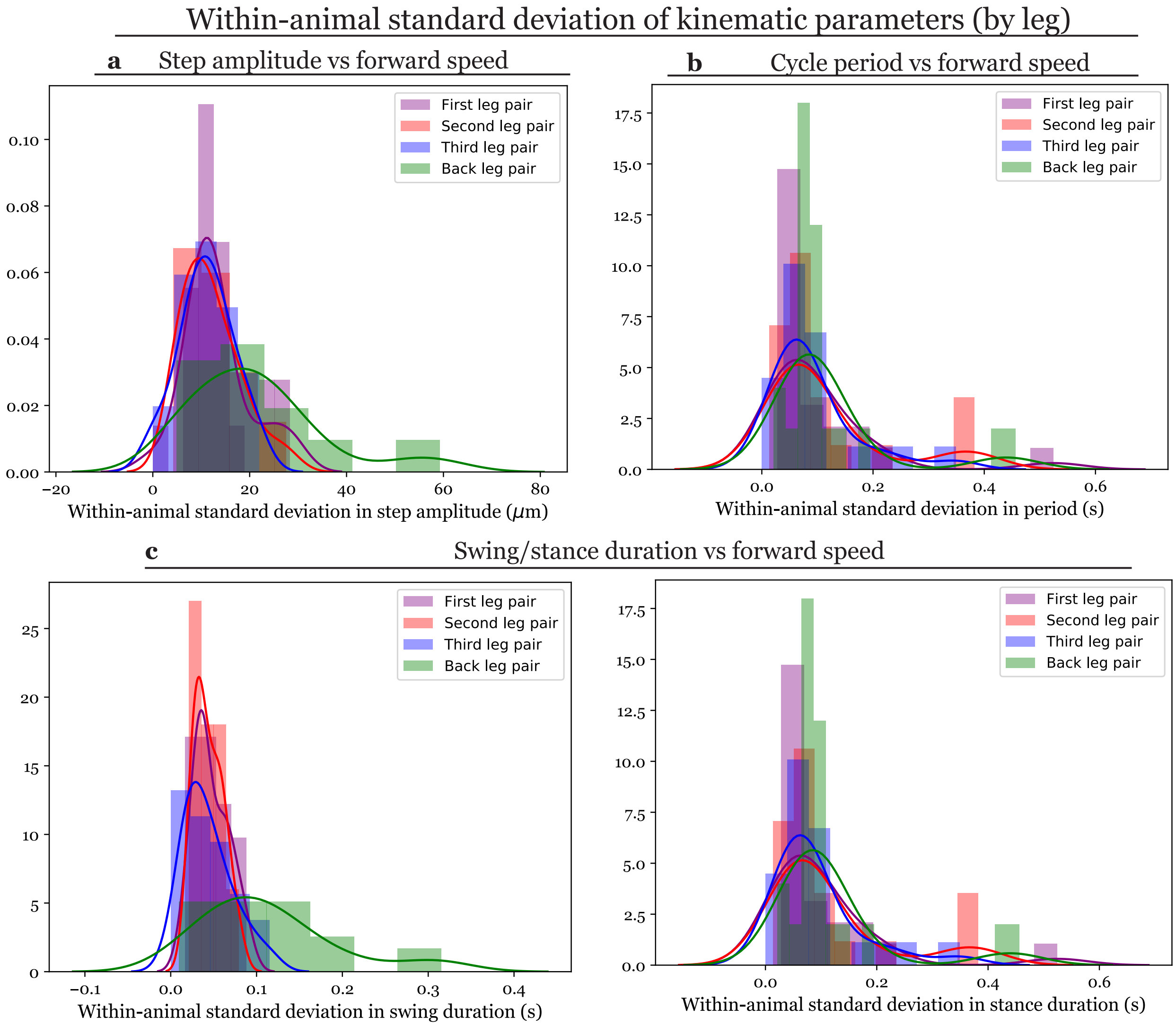

Figure S2: Distribution of within-animal standard deviation for **(a)** step amplitude, **(b)** period, and **(c)** swing and **(d)** stance duration for each leg pair. Data shown for $N = 23$ animals.

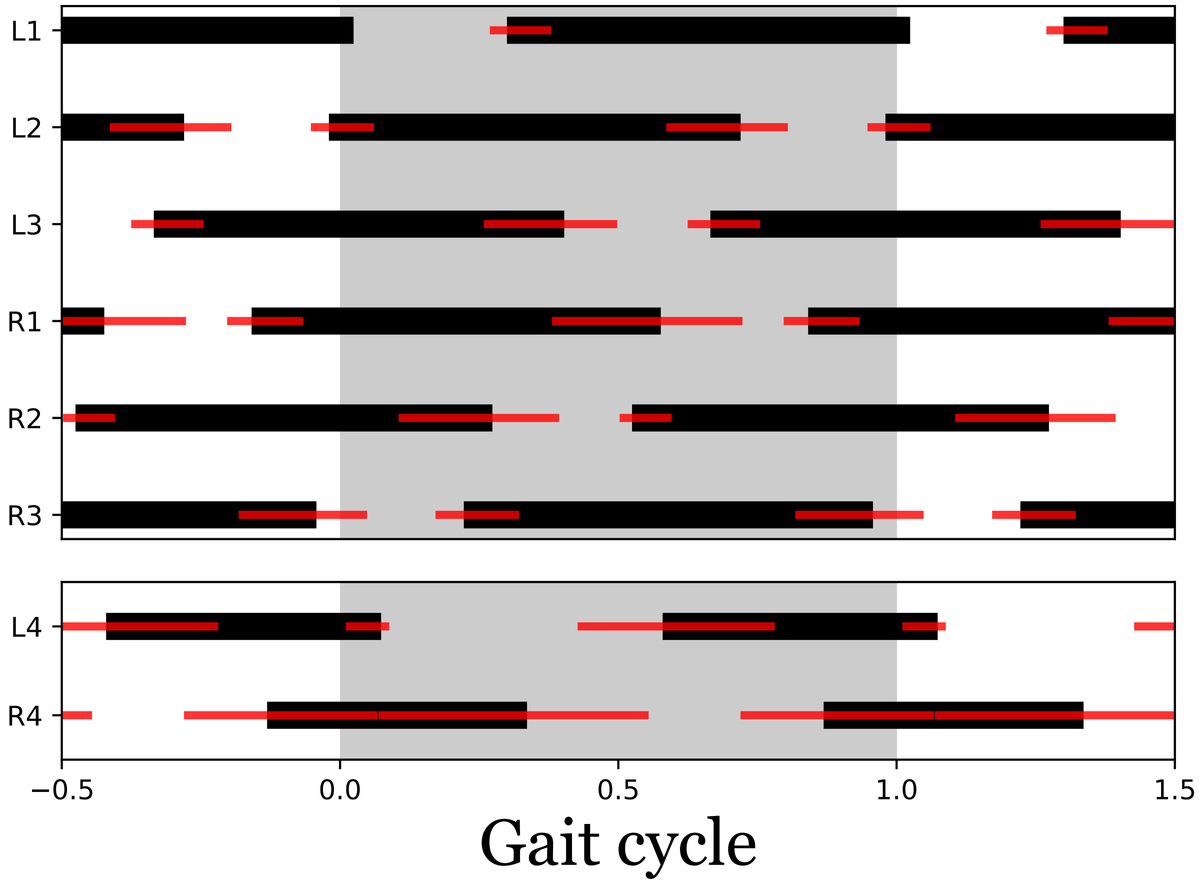

Figure S3: *Averaged gait diagram over all strides (as shown in Fig. 1c) including footfalls of the posterior-most leg pair.* We show the average temporal sequence of ground contacts for legs L1-L4 (left, anterior to posterior) and R1-R4 (right, anterior to posterior). Values are normalized to cycle period of the left front leg L1, shown within the gray shaded region, [0.0 - 1.0]. Extrapolated sequences for previous [-0.5, 0) and subsequent (1, 1.5] periods are shown outside the shaded area.

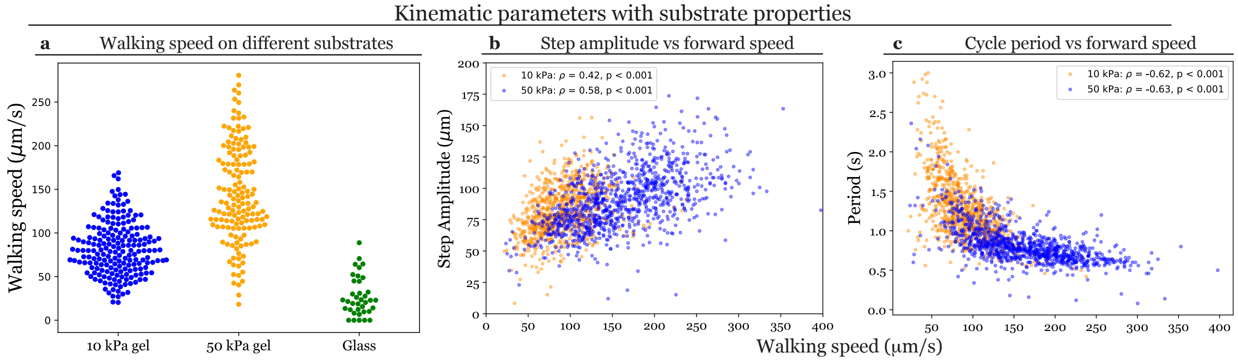

Figure S4: *Comparing walking dynamics on substrates of varying friction and stiffness.* **(a)** Distribution of walking speed on glass and gel of stiffness 50 kPa and 10 kPa. Each point corresponds to the speed of the center of mass (COM) over 1-second windows. **(b)** Step amplitude vs walking speed on gel substrate of stiffnesses 50 kPa and 10 kPa. **(c)** Stride period vs walking speed on gel substrate of stiffnesses 50 kPa and 10 kPa. Each point in **(b)** and **(c)** corresponds to a single stride. Data for all legs are pooled.

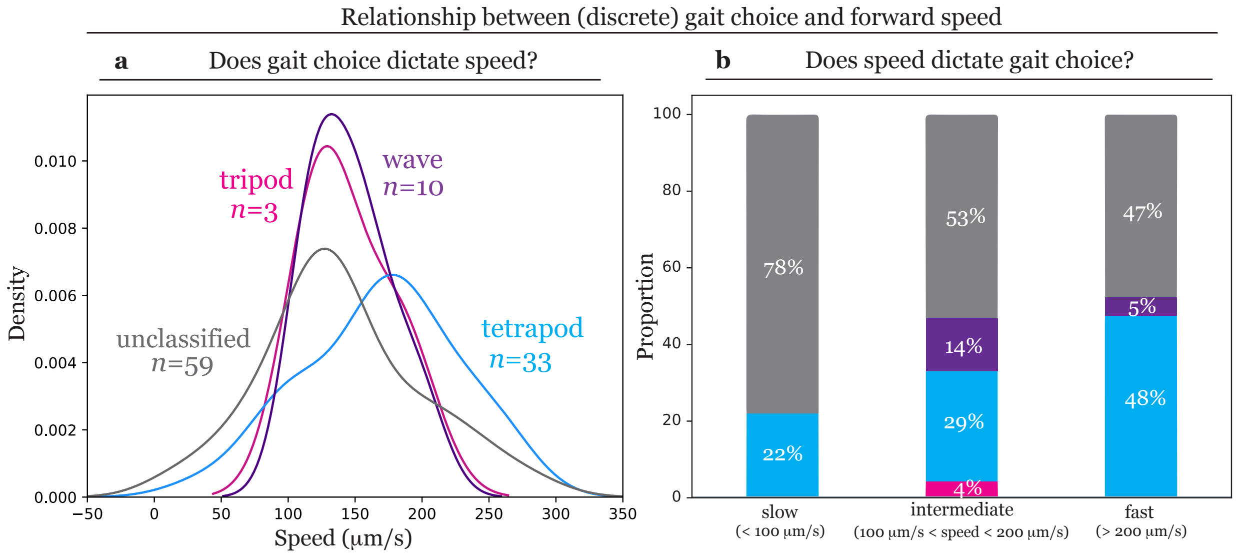

Figure S5: *Relationship between gait choice and forward speed of the center of mass (COM) on 50 kPa substrate.* Distribution of forward speed conditioned upon gait choice **(a)**; and observed frequency of gaits conditioned on forward speed **(b)**. Gait template matching was used to classify each stride into one of three 'canonical' gaits: tripod, tetrapod, or wave gait (see Materials and Methods). The majority of strides (56%) could not be classified; 70% of classified strides showed canonical tetrapod coordination (see main text for more details). Tetrapod gait is observed broadly across all observed speeds (mean ± sd: 160.5 ± 57.8 $\mu$m/s). In **(b)**, each stride was categorized as slow ($v_{\mathrm{COM}} < 100\mu$m/s), medium (100 $\mu$m/s $\leq v_{\mathrm{COM}} \leq$ 200 $\mu$m/s), or fast ($v_{\mathrm{COM}} > 200\mu$m/s). Tetrapod was the most frequently observed canonical stepping pattern in each category. Strides that could not be classified are biased towards lower speeds **(a)**, and noncanonical ('unclassified') strides were more frequently observed within the lowest speed group **(b)**.

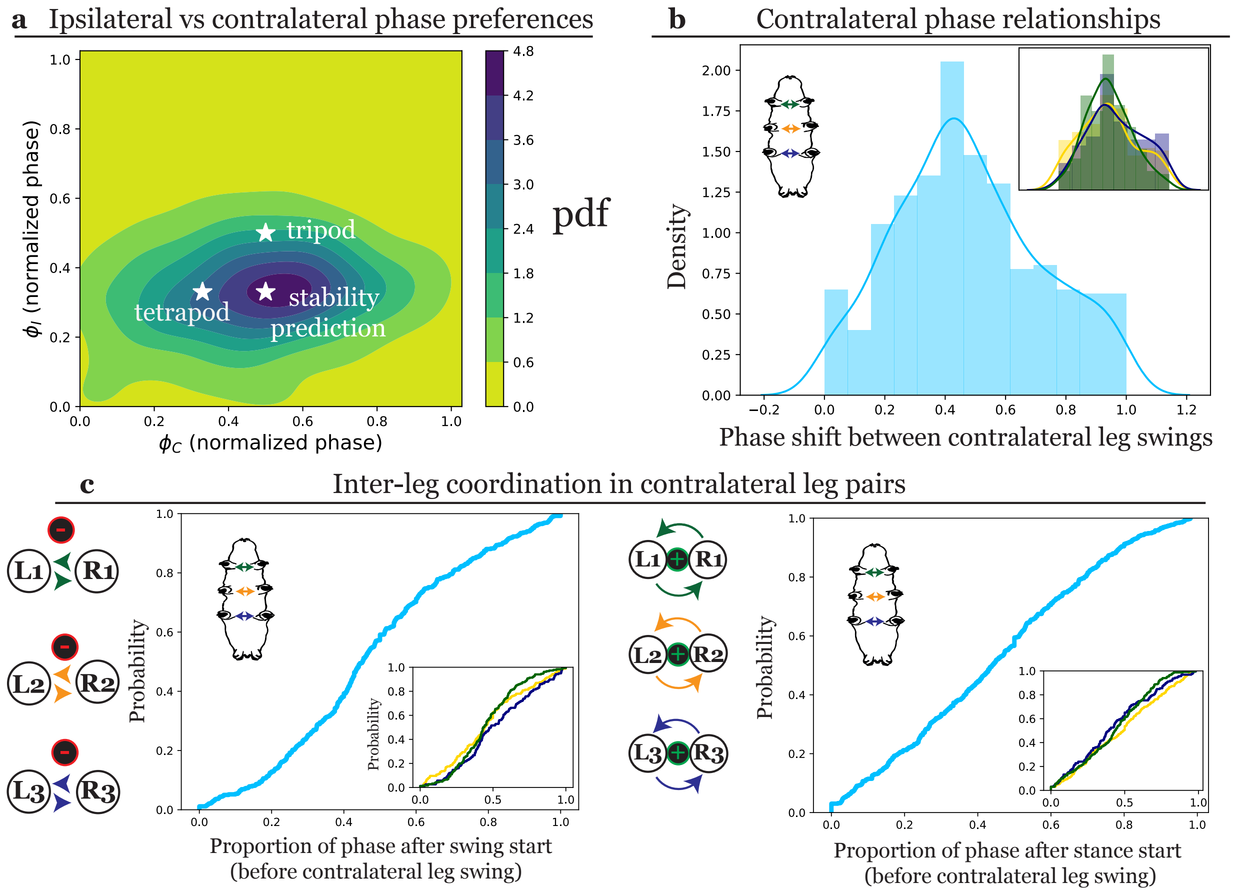

Figure S6: *Temporal coordination between contralateral leg pairs.* Plots represent data pooled from all leg pairs; a total of $n = 654$ strides are shown. **(a)** Joint distribution of the preferred phase offsets between contralateral ($\phi_C$) and ipsilateral ($\phi_I$) leg pairs. Starred points denote phase relationships expected for canonical tetrapod ('tetrapod') or tripod ('tripod') coordination and the predicted optimum from a model for robust static stability ('stability prediction') [7]. **(b)** Histogram and probability density of observed phase shift between contralateral legs show a distribution centered around $\phi_C \approx \frac{1}{2}$. Insets show pairwise inter-leg relationships, with color scheme as demarcated by the arrows on tardigrade schematics. **(c)** Cumulative distribution functions show that leg swings are somewhat suppressed following the swing initiation of the contralateral neighbor (Rule 1 from [1], left). Initiation of swing after the neighboring leg has touched down into stance does not appear to be active between contralateral leg pairs (Rule 2 from [1], right).

[8] Pauli Virtanen, Ralf Gommers, Travis E. Oliphant, Matt Haberland, Tyler Reddy, David Cournapeau, Evgeni Burovski, Pearu Peterson, Warren Weckesser, Jonathan Bright, Stéfan J. van der Walt, Matthew Brett, Joshua Wilson, K. Jarrod Millman, Nikolay Mayorov, Andrew R. J. Nelson, Eric Jones, Robert Kern, Eric Larson, C J Carey, İlhan Polat, Yu Feng, Eric W. Moore, Jake VanderPlas, Denis Laxalde, Josef Perktold, Robert Cimrman, Ian Henriksen, E. A. Quintero, Charles R. Harris, Anne M. Archibald, Antônio H. Ribeiro, Fabian Pedregosa, Paul van Mulbregt, and SciPy 1.0 Contributors. SciPy 1.0: Fundamental Algorithms for Scientific Computing in Python. *Nature Methods*, 17:261–272, 2020.

[9] Michael Waskom and the seaborn development team. mwaskom/seaborn. September 2020.

[10] Anne Wosnitza, Till Bockemühl, Michael Dübbert, Henrike Scholz, and Ansgar Büschges. Inter-leg coordination in the control of walking speed in Drosophila. *Journal of Experimental Biology*, 216(3):480–491, 2013.
